# Cysteine proteases of human hookworm *Necator Americanus as virulence* factors and implications for drug design with anti-heparin and heparin analogs: A bioinformatics study

**DOI:** 10.1101/116921

**Authors:** Arpita Banerjee, Kevin Widmer, Conor R Caffrey, Ruben A. Abagyan

## Abstract

Human hookworm *Necator Americanus (NA)* causes iron deficiency anemia, as the parasite ingests blood from the gastrointestinal tract of its human host. This bioinformatics-based study focuses on eight of the cathepsin B-like cysteine proteases (CPs) of the worm to explore their pathogenic potential. CP1 - CP6, which harbored the active site cysteine residue for enzymatic activity, were relevantly observed to have N-terminal signal peptide for extracellular localization. The secretory CPs could be releasing indigenous worm heparin at the host-pathogen interface for anticoagulation purposes. CP2 and CP3 showed a novel hemoglobinase motif that could be a prerequisite for hemoglobin degradation. CP1 and CP6 shared similar enzymatic-pocket features with cathepsin B and cruzain that cleave high molecular weight kininogen for blood-thinning activity. CP1, CP2, CP3, CP5 and CP6 were predicted to bind heparin, at their C terminal domain, like human cathepsin B and cruzain non-covalently bind heparin to enhance their activity. *NA* CPs’ action in concert with heparin, have implications for anti-heparin and heparin analog design against hookworm infection.

## 1 Introduction

Hookworm infection in humans is a neglected tropical disease that affects over 700 million people worldwide, mostly in the developing countries of the tropical and subtropical regions. ^[1] [2]^ *Necator Americanus (NA)* species of hookworm constitutes the majority of these infections (∼85%) ^[3]^

The clinical manifestation of the disease includes anemia, malnutrition in pregnant women, and cognitive and/or physical development impairment in children ^[4]^. These helminth blood feeders on reaching maturity can feed up to 9ml of blood per day in an infected individual by attaching themselves to the intestinal mucosa of the host, through cutting plates as in *NA* ^[5]^. Iron deficiency anemia is the direct effect of the hookworm’s blood feeding ^[6]^, resulting in other subsidiary consequences of the hookworm disease ^[7] [8]^.

The infective larval stage (L3) worm penetrates into the host skin from soil ^[9] [10]^ and then invades the circulatory system to reach heart and lungs, wherefrom it migrates to alveoli and then to trachea. The parasite eventually reaches gastrointestinal tract as fourth stage larvae (L4) to develop into blood feeding adult stage hookworms ^[1]^

An array of diverse enzymes and molecules in *NA*’s biomolecule repertoire facilitate the pathogen’s survival in the host for up to seven years or longer during the different stages of its lifecycle ^[5]^. The most important therapeutic targets are the enzymes involved in interaction with host and in nutrient acquisition. These are often found in the excretory-secretory (ES) products of the worm. The ES proteins have been shown to engage in crucial functions like tissue degradation for host invasion ^[5] [11]^, fibrinogen degradation ^[12]^ for preventing blood clots, hemoglobin degradation ^[12]^ for nutrient acquisition, and evasion of the host immune system ^[13]^. The enzymes in the ES potpourri of *NA* are not yet completely characterized. However, Brown *et al*, had reported cysteine and serine proteases from ES products to degrade hemoglobin and fibrinogen, and detected the presence of at least two cysteine proteases operating at different pH optima. ^[12]^

The cysteine proteases (CPs) in *NA* form the ninth most gut-expressed abundant gene family ^[14]^ These are most similar to *H.Contortus* (a blood feeding ruminant) CPs that dominate (∼16%) intestinal transcriptome of the barber pole worm ^[15]^, highlighting the importance of pathogenic CPs in host blood degradation. The CPs are synthesized as precursor molecules, where in their folded proenzyme form, a self-inhibitory peptide blocks their catalytic domain. The removal of the inhibitory peptide upon proteolytic action of other peptidases releases the mature cysteine proteases, which are enzymatically active. The active-site clefts of the *NA*-CPs contain the catalytic triad residues: Cysteine, Histidine and Asparagine. Classification of cysteine proteases relies on the sequence homology spanning the catalytic residues ^[16]^. CPs of parasitic organisms are divided into clans CA and CD. The clans are further classified into families in which Cathepsin B-like proteases belong to C1. The *NA*-CPs which are cathepsin B-like, therefore belong to the clan CA C1 family of cysteine proteases, having the typical papain fold comprising of a mostly α-helical amino-terminal known as the L domain, and the antiparallel β strands dominated carboxy-terminal is known as the R domain. The active site for proteolytic degradation is at the interface of these L and R domains ^[17]^. The entire cascade of hemoglobin degradation in hookworm has not been elucidated. However, certain key enzymes have been shown to participate in the degradation pathway. Aspartic protease *NA*-APR1 acts on hemoglobin, whereas metalloprotease MEP1 and cysteine protease CP3 degrade globin fragments ^[18]^. While ingestion and digestion of blood, anticoagulant proteins are secreted to prevent clot formation ^[19] [20] [21]^. Cysteine proteases, from *NA* and from phylogenetically close *H. contortus* have been implicated to have anticoagulation properties ^[12] [22]^. *NA* adopts a number of complementary strategies to evade host procoagulation system ^[21]^, few of which have been elucidated so far.

Taken together, cysteine proteases from different pathogenic organisms perform diverse functions pertaining to blood feeding. This study focuses on eight cysteine proteases viz: CP1, CP2, CP3, CP4, CP4b, CP5, CP6, CP7 in the *NA* genome; of which CP2, CP3, CP4 and CP5 genes are reportedly expressed in abundance in the gut tissue of the adult worm ^[23]^. Only CP3 amongst these has been characterized as a globinase ^[18]^ so far. The other CP genes are suggestive of the involvement of the proteases they encode, in digestive or other functions related to the hookworm’s survival. Despite *NA*-CPs’ importance in the parasite’s physiology, they are under-researched. Not much is known about the function of the individual proteases or about their localization in the ES products of *NA*. This bioinformatics based study on the molecular characterization of the CPs probes into those aspects of the proteases, which could have pathogenic potential such as hemoglobin degradation and blood clot prevention, through the possible assistance of other important small molecules. Several bioinformatics methodologies have been applied viz: sequence-based predictive methods, motif derivation from sequence patterns, homology modeling, docking, and mapping of molecular interactions to elucidate the role of the CPs as possible virulence factors and hence target for therapeutics. The approaches to some of the methods adopted here are in the context of other relevant cysteine proteases.

## 2 Materials and Methods

The following *NA* cysteine protease sequences were retrieved for analyses from Uniprot protein sequence database ^[24]^, (Uniprot ID in parentheses): Necpain or CP1 (Q9U938), CP2 (A1YUM4), CP3 (A1YUM5), CP4 (A1YUM6), CP4b (W2TRZ7), CP5 (A1YUM7), CP6 (W2T0C4) and CP7 (W2SQD9) (the organism code part of the ID is omitted for brevity). The alignments were done in ICM ^[25]^ by BLOSUM62 scoring matrix, with gap opening penalty of 2.40 and gap extension penalty of 0.15. Patterns of relevance and predicted or deciphered sites of functional importance were mapped onto the aligned CP sequences to denote their positions in the protein sequences.

### 2.1 N-terminal-signal pattern and protease localization

The N-terminals of protein sequences often hold clue to protein sorting ^[26]^ which determine the localization of the proteins. CP1-6 harbored pre-sequences at their N-terminals unlike CP7. These pre-sequences were used to derive pattern(s) unique to CP1-6, using PRATT ^[27]^, which were then searched in CP7 by ScanProsite ^[28]^ to see if the latter lacks such pattern(s). The purpose was to delineate the N-terminal signal of CP1-6 for the proteases’ differential localization, as compared to CP7.

The N-terminal signal peptides of cysteine proteases are cleaved off to generate the zymogens ^[17]^. SignalP ^[29]^ was used to predict cleavage sites in the hookworm *NA* CPs to confirm if the N-terminal cleavage products span the signal peptide pre-sequences of the proteases.

The *NA*-CP sequences were submitted for localization prediction to TargetP ^[30]^, iPSORT ^[31]^, TMHMM ^[32]^, LocSigDB ^[33]^, Bacello ^[34]^, Protein Prowler ^[35]^, Cello ^[36]^ and PrediSi ^[37]^ webservers to determine which of the proteases would be prone to secretion. While the algorithms for most of these programs take N-terminal signals into account, Bacello predicts localization on the basis of the information contained in the folded protein and LocSigDB is a signature pattern database derived from proteins whose localization has been confirmed by experiments on bench.

### 2.2 Motif prediction pertaining to hemoglobin degradation

The incompletely elucidated hemoglobin degradation pathway in *NA* describes the role of only CP3 as a globinase, amongst other CPs in the cysteine protease repertoire. In an effort to investigate the involvement of the rest of the *NA* CPs in hemoglobin degradation, cysteine protease sequences from other organisms - known to degrade hemoglobin - were taken along with *NA*-CP3 to derive conserved patterns unique to these proteins. Those organisms, the proteins, and their genbank ^[38]^ accession numbers (in parenthesis) are: *Necator Americanus* CP3 (ABL85237.1), *A. Caninum* CP1 (Q11006), *A. Caninum* CP2 (AIG62903.1), *S.mansoni* CB1 (3QSD_A), *P.falciparum* falcipain2 (AAK06665.1), *P.falciparum* falcipain 3(KOB61544.1), *S.japonicum* Cathepsin B (P43157.1), *H. Contortus* AC3 (Q25032), *H.Contortus* AC4 (Q25031), *P.Westermani* CP1 (AAF21461.1) and *O. ostertagi* CP1 (P25802.3) and *A. Suum* CP (AAB40605.1) Conserved patterns from the aforementioned proteins were derived in PRATT and the motifs were scanned against some other non-hemoglobin degrading proteins in ScanProsite to pinpoint patterns specific to the hemoglobin degrading enzymes. The set of non-hemoglobin degrading organisms and their relevant proteins were: *C. elegans*_CPR3 (AAA98789.1), *C.elegans*_CPR4 (AAA98785.1) and *L. major* cathepsin B (AAB48119.1). Such derived patterns (when found) specific to the hemoglobin degrading enzymes were scanned in the rest of the *NA* CP sequences.

### 2.3 Molecular feature detection for Kallikrein-like activity

Kallikrein-like activity of cleaving high molecular weight kininogen (HMWK) has been reported for cathepsin B and cruzain ^[39] [40]^, which is a cysteine protease of *T.Cruzi* – a parasite that traverses blood capillary vessels as trypomastigotes ^[40] [41]^. Such kininogen-cleaving activity of cruzain and cathepsin B generates bradykinin, a potent vasodilator that releases PGI_2_ - which inhibits platelet activation and degranulation ^[42]^. As *NA* larvae too traces migratory route through blood capillaries of human hosts, the cathepsin B-like cysteine proteases of the human hookworm were scrutinized for molecular features which could be responsible for kallikrein-like activity. This was done by aligning the secretory *NA* CPs with the sequences in the Protein Data Bank (PDB) ^[43]^ deposited structures of human cathepsin B (ID: 1HUC) and cathepsin L-like cruzain, (ID: 2OZ2) – both of which share similar substrate specificity that has been attributed to a critical Glu placed at their S2 subsites, in the protease structures ^[44]^. The sequences were aligned in ICM with the same parameters as mentioned before. The distances between the C-alpha atoms of relevant S2 subsite residues within the enzymatic cleft of the protease structures were measured in ICM for estimating the possible kininogen-cleaving role of the crucial residues (if any) in the *NA* CPs.

### 2.4 Heparin-binding: motif, domain, and docking simulation

Glycosylation with glycosaminoglycans (GAGs) like heparin is an important post-translational modification in cysteine proteases of many pathogenic organisms ^[16]^. The CPs of *NA* hookworm were therefore scanned for sequence motifs that could undergo such modification. Scanprosite was used for searching putative glycosylation motifs, which could be the covalent attachment sites for pathogenic glycosaminoglycan (indigenous to the worm) like heparin. Such sites were searched to explore the possibility of (worm) heparin, with anticoagulation properties, being transported out to extracellular matrix while attached to the secretory *NA* CPs (if any), for impairing host coagulation system.

Importantly, glycosaminoglycan molecules remain displayed on host proteoglycans ^[45]^ and are probably encountered for recognition by the worm ES products at the host-pathogen interface. The sequences were subjected to query by ProDom ^[46]^, a protein domain family database derived from Uniprot knowledgebase, for finding any heparin-binding protein domains on the *NA* CPs; as such sites would be predisposed to non-covalently bind heparin that remain covalently attached to host proteins. Structure-wise, PDB ligands SGN and IDS, which constitute the GAG tetramer in the heparin-bound structure 5D65, were used to construct a frequency table in ICM, from the residues within 4Å of the ligands, in all the PDB-deposited structures harboring the SGN and IDS units. This was done to figure out the residue-environmental preference for such heparin ligands, in order to confirm the residue-profile of the sequence-derived heparin-binding motif.

*NA* CP - heparin dockings were then undertaken to explore the feasibility of such recognition of host proteins, through GAG interaction, by the hookworm’s CPs. Homology models of the *NA* cysteine proteases were built to carry out the CP – heparin dockings, due to lack of experimental three-dimensional structure of the CPs. BLAST search for finding homology model templates was performed against PDB. 3QSD - mature CathepsinB1 enzyme from *Schistosoma Mansoni* - was chosen from the search results for building models as it aligned well at the active site and had a resolution of 1.3 Å. Also, this structure had co-ordinates for the two occluding loop residues near the active site, which have been designated crucial for the exopeptidase activity for this class of cathepsin B-like proteases ^[47]^. Homology models were built within the internal co-ordinates mechanics protocol of ICM software. The sequence alignment between the template and the model sequence was generated by using BLOSUM62 matrix, with gap opening penalty of 2.40 and gap extension penalty of 0.15. Further, for generating reliable models, the alignment around the active site was edited wherever necessary, according to conservation propensity of residues, and for modeling the occluding loop residues. Loops were sampled for the alignment gaps where the template did not have co-ordinates for the model sequence. The loop refinement parameters were used according to the default settings of the procedure. Acceptance ratio during the simulation was 1.25. The *NA-*CP model structures were then built within the full refinement module of the software. The qualities of the homology models were checked using PROCHECK^[48]^ which showed 100% of the residues from most of the CPs to lie within the allowed regions of the Ramachandran plot. CP2 and CP5 were the exceptions, which had 99.5% of residues in the allowed regions.

The co-ordinates of heparin tetrasachharide were retrieved from its complex deposited in PDB (ID: 5D65) in SDF format, with alternating units of N, O6-disulfo-glucosamine (PDB ID: SGN) and 2-O-sulfo-alpha-L-idopyranuronic acid (PDB ID: IDS).

The CP homology models were converted to ICM formatted receptors for docking the heparin molecule. The sequence stretch of the CPs encompassing the predicted fibronectin domain, which can putatively bind heparin like molecules ^[49]^, was selected for docking. The receptor maps were generated with grid step of 0.50. The dockings for non-covalent binding were performed with a thoroughness level of 3, with the generation of three initial ligand conformations for each simulation.

The heparin-bound *NA* CP models were rendered in electrostatic surface representation by ICM, where the potential scale was set to 5.0 along with the assignment of simple charges, for the purpose of viewing the electrostatics of the protein sites that were occupied by the negatively charged heparin.

## 3 Results and Discussion

### 3.1 N-terminal signal for differential localization of proteases

The *NA*-CP alignment showed that the CP7 lacked the N-terminal signal pre-sequence present in the other proteases (**Figure1**). The PRATT derived motif unique to CP1-6’s pre-sequence was M-x(4,5)-L. CP7’s N-terminal however had a lysosomal targeting pattern [DE]x{3}L[LI], according to LocSigDB. The specific residues in the pre-sequences are summarized in **Table1**, along with the lysosome targeting peptide in CP7.

SignalP-derived cleavage sites predicted the length and the peptide sequence for the signals contained in the pre-sequences of CP1-6. **Figure 1** shows the positioning of the cleavage sites, where the signal peptide would be cleaved off to release the zymogens. The lengths of these signal peptides across the *NA* CP1-6 were approximately the same. CP7 did not show any such cleavage site at equivalent position to the other proteases, confirming its lack of similar N-terminal signal, re-emphasizing that this protease gets synthesized without the signal for extracellular localization, unlike the other CPs.

**Figure 1.**
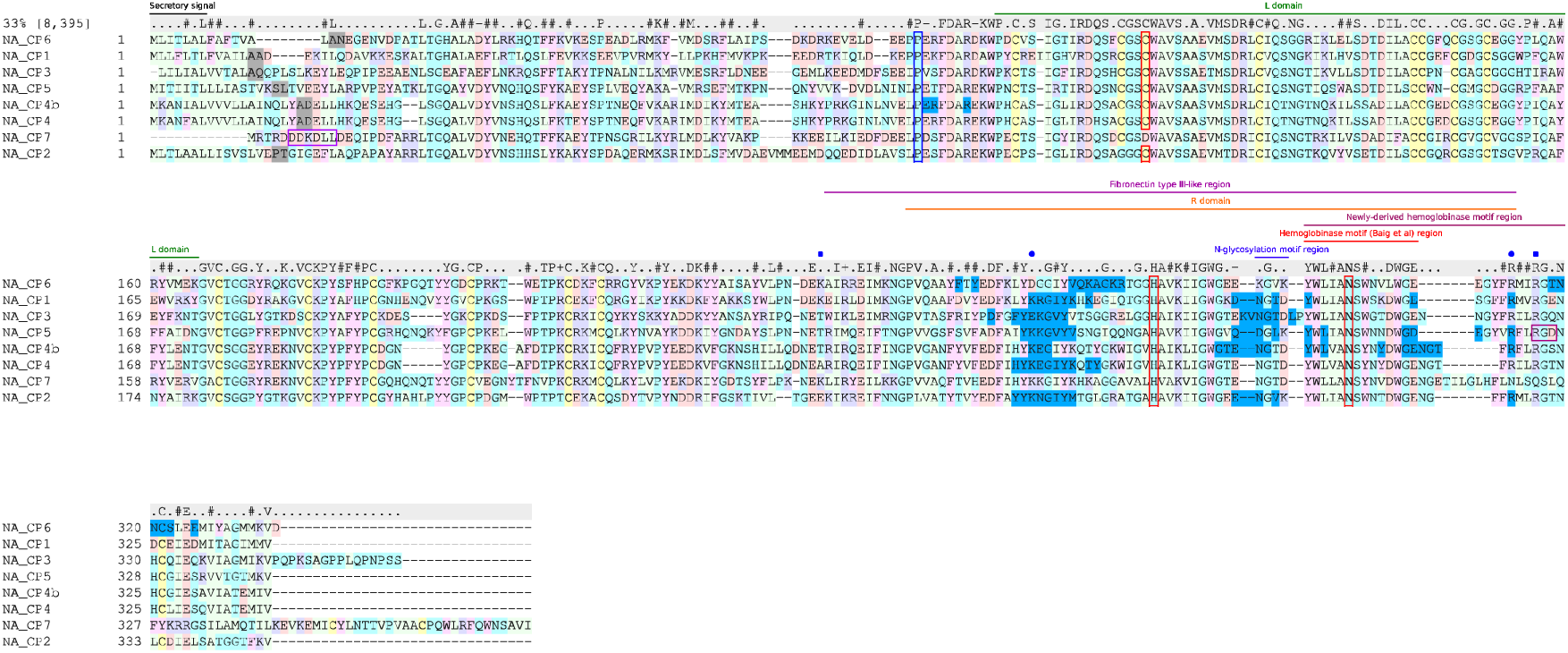
Sequence alignment of the *NA* CPs with annotated regions of functional importance. The residues flanking the N-terminal-signal cleavage sites are shaded in grey. The lysosome-targeting peptide of *NA* CP7 is boxed. The begin position of the mature proteases is boxed in blue and the active-site catalytic triad residues are within red box. The derived hemoglobinase motif (region indicated in dark red) appears in *NA* CP2 and CP3. The RGD motif signature in CP5 for fibronectin type III-like domain is boxed. The residues in contact with the docked heparin molecule are shaded in blue. The K and R residues contacting heparin in most of the docked complexes are indicated by blue spheres to denote their positions with respect to the heparin-binding residues K157 and R234 of human cathepsin B (Costa *et al*, 2010) that has been marked by blue squares.

The consensus from the localization prediction methods deemed CP1-6 to be secretory proteins, having extracellular localization. CP7, lacking the N-terminal pre-sequence, was predicted to be non-secretory with its lysosome targeting signal peptide (**Table 1**).

**Table 1.**
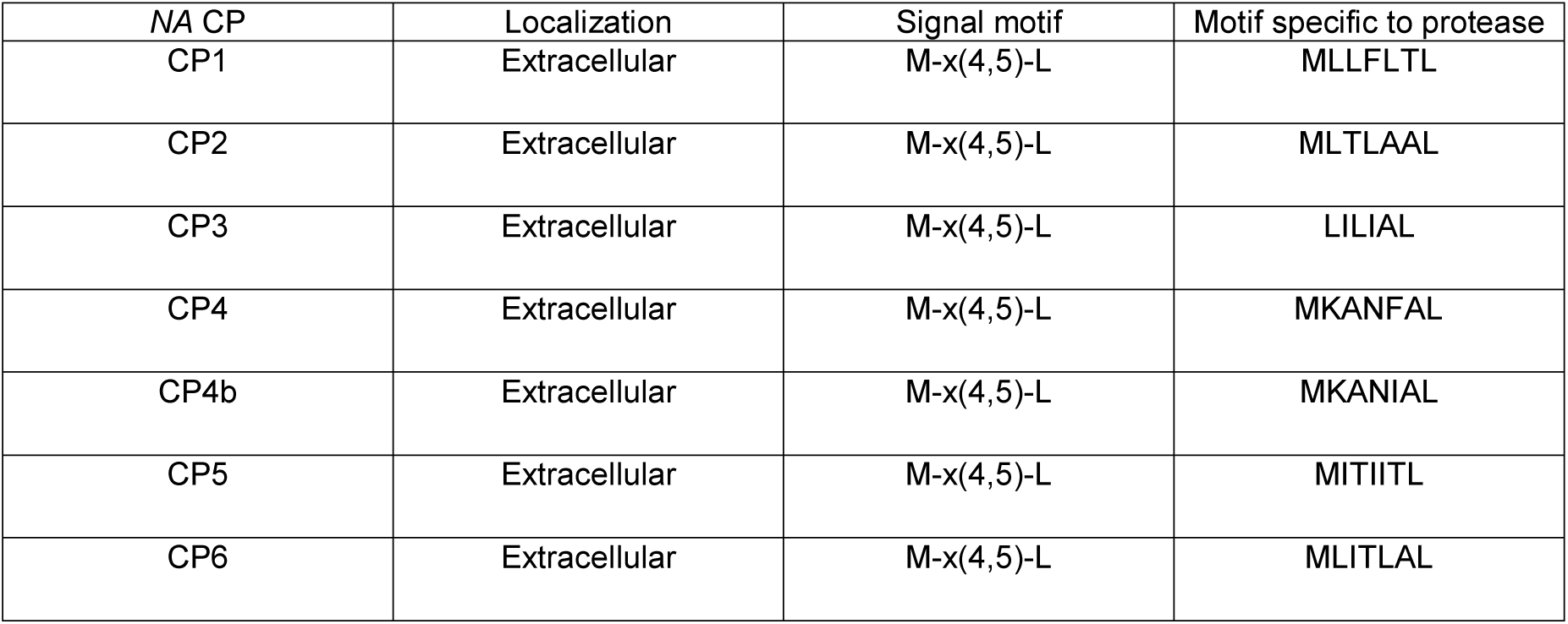
Localization of the *NA* cysteine proteases, and the corresponding motifs pertaining to the N-terminal signals.

| NA CP | Localization | Signal motif | Motif specific to protease |
| --- | --- | --- | --- |
| CP1 | Extracellular | M-x(4,5)-L | MLLFLTL |
| CP2 | Extracellular | M-x(4,5)-L | MLTLAAL |
| CP3 | Extracellular | M-x(4,5)-L | LILIAL |
| CP4 | Extracellular | M-x(4,5)-L | MKANFAL |
| CP4b | Extracellular | M-x(4,5)-L | MKANIAL |
| CP5 | Extracellular | M-x(4,5)-L | MITIITL |
| CP6 | Extracellular | M-x(4,5)-L | MLITLAL |

The conserved hydrophobic M-x(4,5)-L sequence pattern in the *NA* CP1-6 are presumably signal peptides for extracellular localization. N-terminal signals are extremely degenerate across various proteins and the reported motif could be part of the unique signal peptide for these proteins.

CP1-6, being predictively secretory, could be available for important host-pathogen interactions. CP3 amongst these has been implicated to be present in the gut of an adult worm at the host-pathogen interface ^[23]^, and has been shown to be involved in the hemoglobin degradation pathway ^[18]^. Therefore, CP1, CP2, CP4, CP4b, CP5 and CP6 could be expected to perform similar or other functions for the hookworm’s survival. Whereas, CP7 that lacks the active site cysteine residue (and can not degrade other proteins) was predicted to reside in lysosome for unknown purposes.

### 3.2 Novel hemoglobinase motif with implicated hemoglobin-degrading functionality

The hemoglobinase motif Y-[WY]-[IL]-[IV]-x-N-S-W-x-[DEGNQST]-[DGQ]-W-G-E-x(1,2)-G-x-[FI]-[NR]-[FILM]-x(2)-[DG]-x-[DGNS] was derived from the hemoglobin degrading cysteine proteases of the following blood feeders viz: *NA* (CP3), *A. Caninum, S.mansoni, S.japonicum, H. Contortus, O. ostertagi, P.falciparum, P.westermani* and *A. Suum*. The pattern detected here is longer than the previously reported hemoglobinase motif Y-W-[IL]-[IV]–A-N-SW-X–X–D-W-G-E motif ^[50]^. See comparison in **Figure 1**. The newly derived motif was absent in the cysteine proteases of the non-blood feeders viz: *C. elegans* and *L. major*. The observations from this study are similar to the previous study in the context of the presence of the motif in blood feeding proteases and its absence in the non-blood feeding proteases. The derived motif when searched in the *NA*-CPs (excluding CP3 of the training dataset) was detected only in CP2.

The cysteine proteases from the *NA* ES products of the parasite had been demonstrated to cleave hemoglobin ^[12]^. However, only CP3 in the worm has been characterized to have such role. The molecular features pertaining to such function in *NA* CPs had not been researched. In this study, a specific sequence pattern generic to hemoglobin degrading enzymes was sought, without emphasis on a particular family of cysteine proteases. Hemoglobin-cleaving non-cathepsin B enzymes; falcipain2 and falcipain3 from *P.falciparum* and CP1 of *P.Westermani*, were therefore included in the training dataset, which had not been taken into account by Baig *et al*. Despite adopting a different methodology (mentioned in materials and methods) from the previous study for deriving the motif, a pattern unique to the blood degrading enzymes emerged. Working with a longer motif, as reported here, is advantageous in terms of avoiding false positives. CP2 was the only protein in the repertoire of the *NA* cysteine proteases to have the motif, other than the already established globinase CP3 ^[18]^. This observation is suggestive of CP2’s involvement in hemoglobin degradation, along with CP3.

### 3.3 Kallikrein-like activity of cleaving kininogen for implicated vasodilation and anticoagulation

*NA* CP1 and CP6 were observed to have an aspartic acid and a glutamic acid respectively, near the S2 subsite of their enzymatic pocket - similar to human cathepsin B’s and cathepsin L-like cruzain’s critical Glu, **(Figure 2),** which forms an important specificity determinant for substrates having Arg or Phe ^[44]^. HMWK cleavage by cathepsin B and cruzain generates vasoactive bradykinin ^[39] [40]^, which dilates blood vessels and eventually thwarts the blood coagulation cascade. *NA* CP1 and CP6, due to their similarity in enzymatic-pocket molecular feature with cruzain and human cathepsin B, are likely to bind human HMWK as their substrate in the region Leu^373^-Gly-Met-Ile-Ser-Leu-Met-Lys-Arg-Pro-Pro-Gly-Phe-Ser-Pro-Phe-Arg-Ser-Ser-Arg-Ile^393^ of the kininogen; the same region that cruzain cleaves between Met^379^-Lys^380^ and Arg^389^-Ser^390^ to generate the vasodilator Lys - bradykinin ^[40]^. With the catalytic cysteine residue of the cysteine protease enzymes located between S1’ and S1 subsites, where the peptide bond hydrolysis occurs, the cleavage sites of HMWK indicate that Arg^381^ and Phe^388^ of the kininogen are likely to get accommodated at the S2 subsite of cruzain. The side chain of the Glu207 at its S2 site could form salt-bridge interaction with the charged Arg^381^ of the substrate. Next, when the hydrophobic Phe388 of the kininogen occupies S2 subsite, the aforementioned critical glutamate of the enzyme could turn away to face the solvent. Therefore, the proteolytic action on HMWK by *NA*-CP1 and CP6 - which possess acidic Asp246 and Glu245 near their S2 pockets - could possibly release vasoactive peptides.

**Figure 2.**
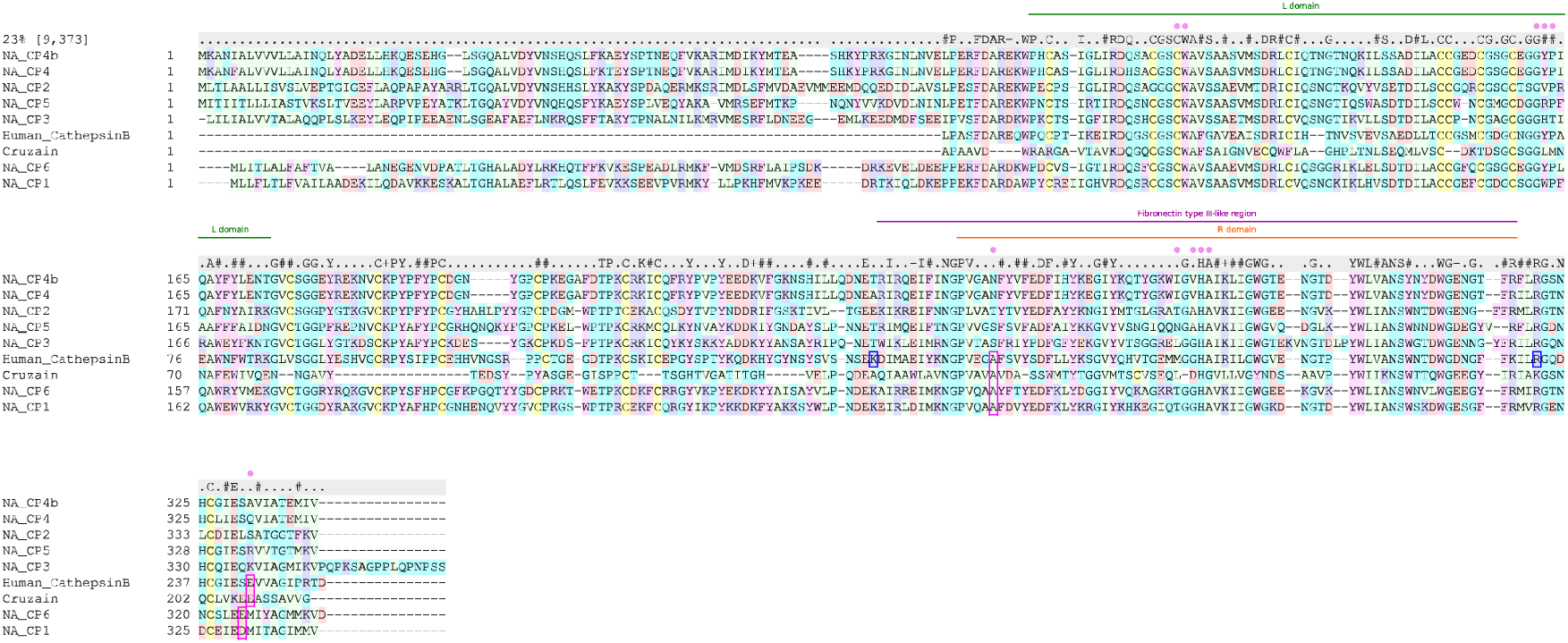
Sequence alignment of the secretory *NA* CPs with cruzain and human cathepsin B – both of which bind heparin and cleave HMWK. Regions of common relevance are annotated. Pink spheres indicate the S2 pocket residues derived from the inhibitor-bound cruzain structure. The conserved Ala and the crucial acidic residues shared between CP1, CP6, cruzain and human cathepsin B, in and around the S2 subsite, are boxed. The K157 and R234 residues hypothesized to be crucial for binding heparin in human cathepsin B (Costa et al, 2010) are boxed in blue.

The enzymatic pocket of cathepsin B, cruzain, *NA* CP1 and CP6 shared an alanine at their S2 subsite (**Figure 2**). Cruzain’s Ala137 and human cathepsin B’s Ala172 at their S2 subsites, located near their critical Glu residue, corresponded with Ala175 of CP1 and Ala174 of CP6. The distances between the C-alpha atoms of this conserved Ala and the critical acidic Asp/Glu residues of the relevant enzymes are listed in **Supplementary Table 1**. Such measurements were made in order to get an estimate of the distance of the critical Asp/Glu from the active site cleft, given the lack of substrate/ligand in the structures (except cruzain). The Ala and Asp/Glu around the S2 site of *NA* CP1 and CP6 were spaced further away compared to the distance between the relevant Ala and Glu in cruzain and human cathepsin B. However, in the dynamic *NA* CP proteins, the side chains of the charged acidic residues could probably move closer (more so Glu245 of CP6, because of its longer side chain) to make the salt-bridge interaction with charged Arg^381^ of HMWK substrate.

Human plasma kallikrein (pKal) is a component of the anticoagulation pathway that cleaves HMWK - a protein involved in blood coagulation and fibrinolysis - to produce a potent vasodilator bradykinin ^[42]^ that stimulates prostacyclin (PGI_2_) release from endothelial cells. PGI_2_ in turn is an inhibitor of platelet activation and degranulation pathway ^[42]^. Emulation of plasma kallikrein activity has been reported for cruzain ^[40] [41]^ – a cysteine protease from *T.Cruzi* that traverses blood capillary vessels. Such function has been suggested also for SmSP1 ^[51]^ - serine protease of blood fluke *S.mansoni*, as kallikrein-like activity had been reported for a serine protease from *S.mansoni* ^[52] [53]^. The blood stream navigating physiology of the mentioned pathogens perhaps explains their need for cleaving HMWK to generate bradykinin for vasodilation and blood clot prevention.

Extracellular cathepsin B has also been implicated to participate in kininogenase activity ^[39]^ and has been shown to impair coagulation and fibrinolysis under pathological conditions ^[54]^. Cathepsin B-like extracellular CP1 and CP6 of the blood stream navigating *NA* could possibly generate bradykinin by the dint of their critically positioned Asp246 and Glu245. The hookworm might be exploiting this component of the anticoagulation pathway (amongst others) to evade host blood clots for easing migration.

### 3.4 Heparin binding for transportation of indigenous worm glycosaminoglycans, and for molecular recognition of host proteins

Asn glycosylation ‘NGTD’ motifs were detected in *NA* CP1, CP3, CP4, CP4b and CP7, as per prosite entry PS00001. The glycosylation site region in the alignment (**Figure1**) showed CP2 to retain at least the Asn residue for N-glycosylation in its ‘NGVK’ stretch. The prediction of N-glycosylation motifs within the predicted fibronectin type III domain region of the *NA* CPs hint that these proteases can have GAG molecules covalently attached to these sites as post-translational modifications. Such modifications could include heparin, which are reported to occupy N-glycosylation sites, and even sites without the Asn attachment point ^[55]^. N-glycosylation sites harboring *NA*, CP1, CP2, CP3, CP4 and CP4b, could be transporting the hookworm’s indigenous heparin (if present) to the host-pathogen interface. CP5 and CP6, which did not show any N-motif in the region, could also be glycosylated with heparin. Heparin when released at the interface, by deglycosylation of the secreted *NA* CPs, could possibly assist in evading blood coagulation (by serving as co-factors to antithrombin - an inhibitor of thrombin and other coagulation factors in the blood plasma), as has been implicated for indigenous worm heparin/heparan sulfate from the tegumental surface of blood fluke *S.mansoni* ^[51]^.

CP2 and CP5 sequences were predicted by ProDom to have fibronectin domain type III (entry: PDC9H7K4), which is known to harbor N-linked and O-linked glycosylation sites ^[49]^ and has been shown to bind heparin ^[56]^. The N-glycosylation sites mentioned earlier were within the region spanning this predicted fibronectin III domain in the *NA* CPs. The region did not have any cysteine residues in any of the proteases for disulfide bond formation, which is again a feature of type III repeat ^[49]^. The domain region was 74 amino acids in length. CP3 having the longest sequence had 79 residues. CP5 showed the RGD motif- bordering the predicted fibronectin region (See alignment in **Figure 1**), which is another signature for type III_10_ repeat ^[49]^. With CP5 as reference, the fibronectin domain-like region in the rest of the secretory CPs showed sequence identity within 50.64% and 55.26%, and the sequence similarity ranged from 65.11% to 78.64%. CP2 and CP5 were most similar in the region. Relevantly, CP7 that is apparently not secreted and would not require binding host heparin from functional point of view, shared sequence identity of only 40.74% and sequence similarity of 55.15% with CP5 in the fibronectin-III like region. (**Supplementary Table 2**).

The R domains (overlapping with fibronectin III, **Figure 2**) of the modeled-secretory proteases did not exactly resemble the beta sandwich fold typical of fibronectin III. Instead, they resembled the antiparallel beta strand and loop dominated R domains of *heparin-binding* human cathepsin B and cruzain. The clustering map generated from the sequence alignment of the R domains of the secretory *NA* CPs (cathepsin B-like) with those of the human cathepsin B and cruzain, placed human cathepsin B closer to the CPs. Cruzain, which is cathepsin L-like understandably, shared weaker identities with the hookworm proteases. **Figure 3A.**

**Figure 3.**
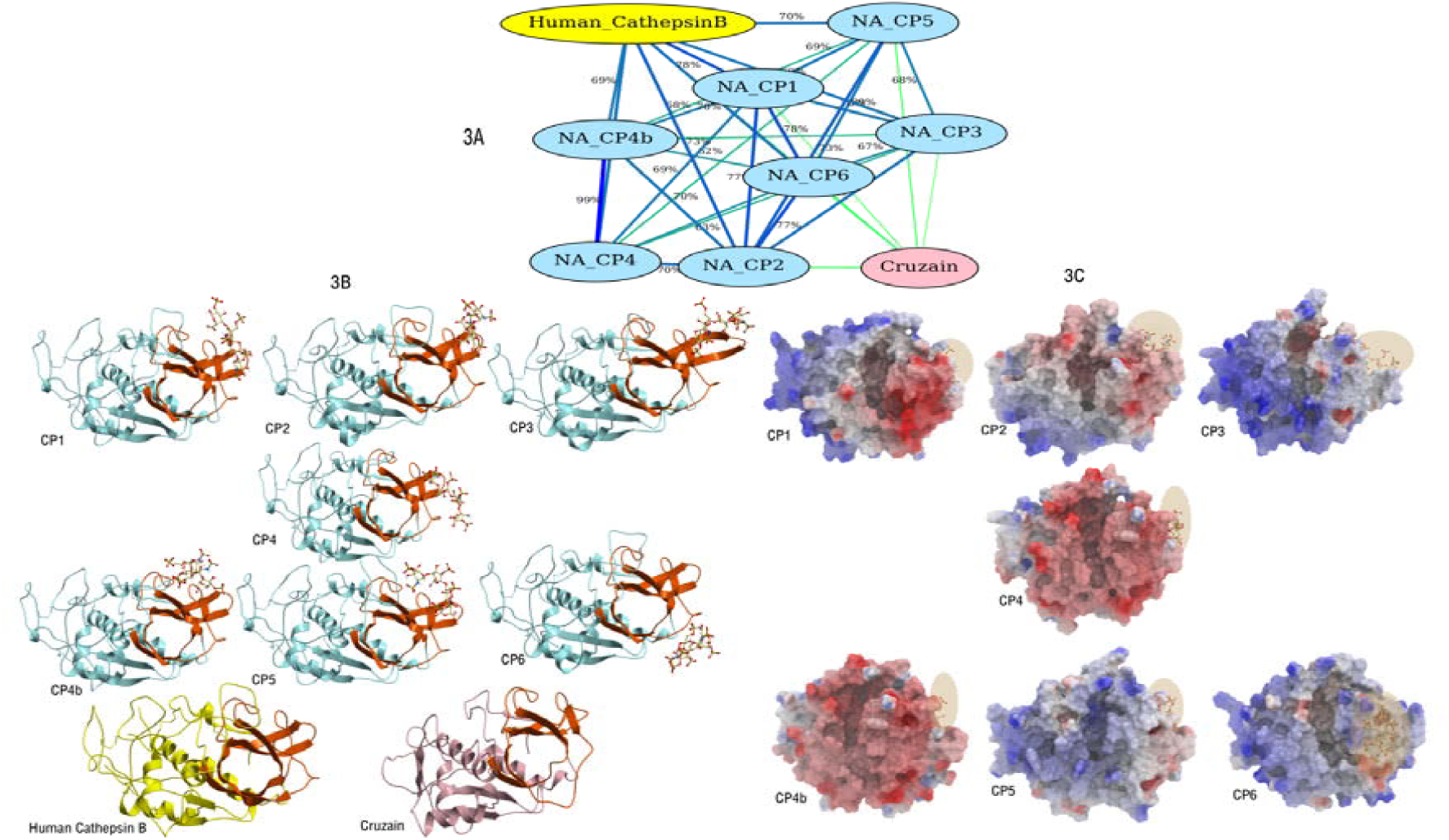
**A.** Distance-wise sequence clustering maps of the R-domains of secretory *NA* CPs, Cruzain, and Human cathepsin B. **3B.** The R domains at the C-terminus (orange) of the secretory *NA*-CP homology models, compared to the structurally similar R domains of Cruzain (PDB ID: 2OZ2) and human Cathepsin B (PDB ID: 1HUC) - implicated to bind heparin. The antiparallel beta strand and loop dominated domain overlaps with the predicted fibronectin type III-like region – where heparin was docked. Heparin (represented in stick) mostly made contacts with the loops of the highlighted domain. **3C.** Heparin-docked homology models of the secretory NA-CPs in electrostatic surface representation. The active sites of the cysteine proteases appear as clefts at the front. The locations of the heparin (represented in stick) that bind away from the active site are highlighted within ovals.

The sequence-structure profile of the secretory CPs implied that they would be predisposed to bind heparin at the predicted fibronectin III-like domain. Heparin on being docked at the putative *heparin-binding* site of the cathepsin B-like secretory CPs showed the best scored conformation to bind surface loops **Figure 3B**, away from the enzymatic cleft, at a site similar to what has been reported in an earlier docking/MD simulation study on human cathepsin B - heparin interaction ^[57]^. The binding site residues within 4Å of the ligand are underscored in the sequence alignment (**Figure 1)**. Barring CP6, heparin occupied almost similar sites in the other CP homology models **(Figure 3B)**. The positioning of heparin was slightly shifted in CP6, albeit away from the active site (**Figure 3C**). The negative sulfate groups in the glycosaminoglycan molecule mostly interacted with basic/neutral residues at the binding site. Some of the heparin-contacts of the CPs showed the sequence patterns: BBX, XBBX, BXB, and BXXBB, which were entirely or partially in conformity with previously reported heparin-binding motifs ^[58] [59] [60] [61]^, where B is a basic residue and X is any amino acid. The overall electrostatics (**Figure 3C**) at the heparin-bound sites of the CPs varied from predominantly neutral to positive (except CP2, CP4 and CP4b), depending on the site-residues and long-range electrostatic effects from residues beyond. The highly negative binding-site electrostatics of CP4 and CP4b (which shared least sequence identity/similarity with CP5 in the fibronectin-III like region) contributed to unfavorable docking scores for these proteases. The rest of the CPs showed good scores for binding heparin, which were dominated by favorable van-der Waals interactions, with the contact residues being mostly basic and hydrophobic. **Table 2** summarizes the scores, contact residues, sequence patterns, and H-bonding interactions of the highest scored conformations of heparin. The crucial interactions made by human cathepsin B’s basic residues Lys157 and Arg234 for binding the heparin ^[57]^, highlighted from structural studies by Costa *et al*, mapped close to this study’s heparin-interacting Lys and Arg residues in most of the *NA*-CPs (derived from the alignment of human cathepsin B with *NA* CPs: **Figure 2**). This is the closest comparison (**Figure 1**), which could be drawn, with no structures of cathepsin B – heparin complex available in PDB.

**Table 2.**
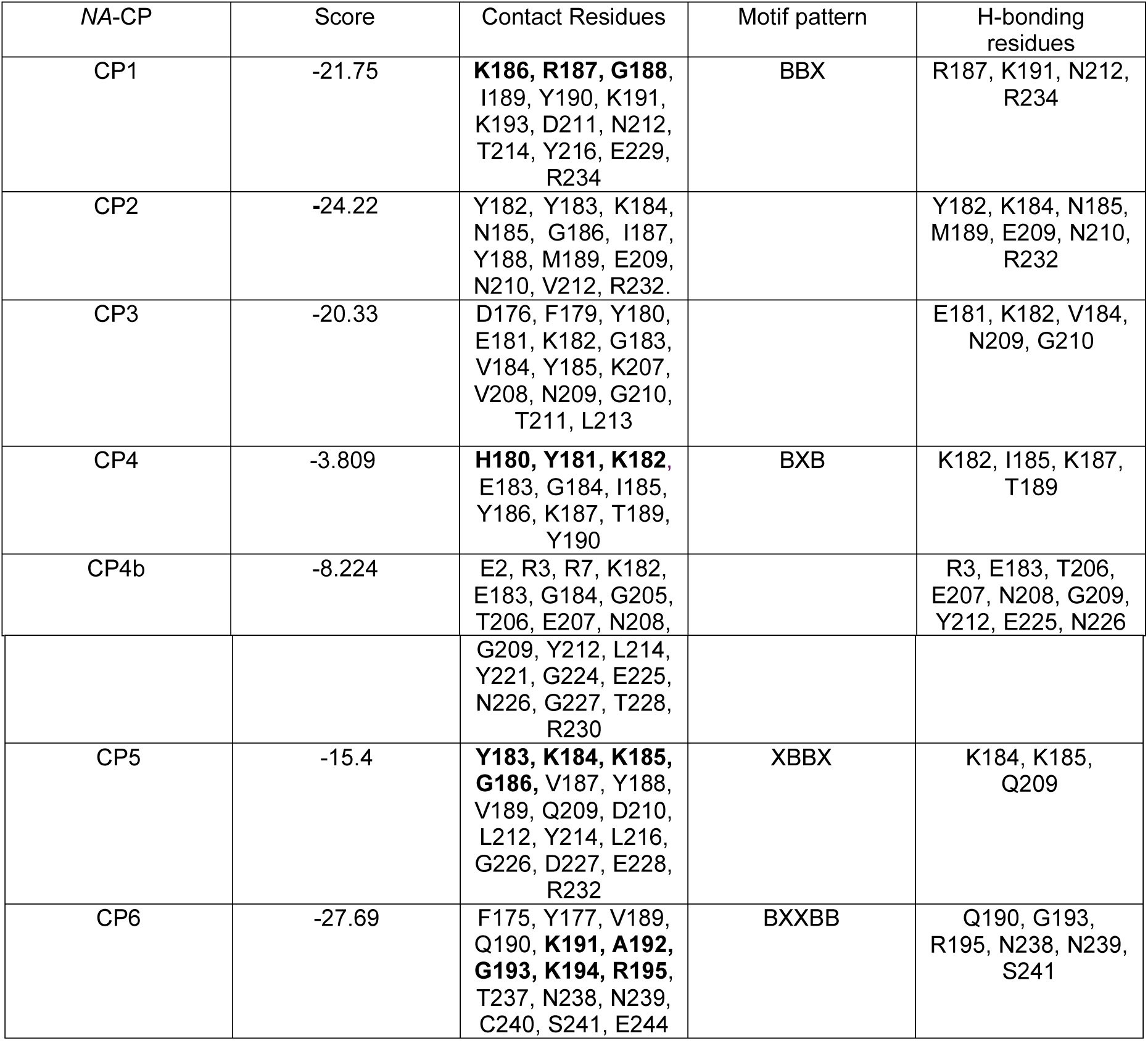
Docking scores, contact residues, sequence motif patterns, and H-bonding interactions of the highest scored conformations of the heparin molecule docked onto the *NA* cysteine proteases. The residue numbering pertains to the mature proteases.

| NA-CP | Score | Contact Residues | Motif pattern | H-bonding residues |
| --- | --- | --- | --- | --- |
| CP1 | -21.75 | **K186, R187, G188**, I189, Y190, K191, K193, D211, N212, T214, Y216, E229, R234 | BBX | R187, K191, N212, R234 |
| CP2 | -24.22 | Y182, Y183, K184, N185, G186, I187, Y188, M189, E209, N210, V212, R232. |  | Y182, K184, N185, M189, E209, N210, R232 |
| CP3 | -20.33 | D176, F179, Y180, E181, K182, G183, V184, Y185, K207, V208, N209, G210, T211, L213 |  | E181, K182, V184, N209, G210 |
| CP4 | -3.809 | **H180, Y181, K182**, E183, G184, I185, Y186, K187, T189, Y190 | BXB | K182, I185, K187, T189 |
| CP4b | -8.224 | E2, R3, R7, K182, E183, G184, G205, T206, E207, N208, |  | R3, E183, T206, E207, N208, G209, Y212, E225, N226 |
|  |  | G209, Y212, L214, Y221, G224, E225, N226, G227, T228, R230 |  |  |
| CP5 | -15.4 | **Y183, K184, K185, G186**, V187, Y188, V189, Q209, D210, L212, Y214, L216, G226, D227, E228, R232 | XBBX | K184, K185, Q209 |
| CP6 | -27.69 | F175, Y177, V189, Q190, **K191, A192, G193, K194, R195**, T237, N238, N239, C240, S241, E244 | BXXBB | Q190, G193, R195, N238, N239, S241 |

Fibronectin can non-covalently bind a number of biologically important ligands that includes glycosaminoglycan such as heparin or heparan sulfate, which are present on the surfaces of extracellular proteoglycans ^[49]^. The molecule’s type II domain forms the primary binding site for heparin. However, fibronectin fragments with type III_7-10_ and type III_10-12_ repeats have been shown to bind heparin ^[56]^. In the cathepsin B-like *NA*-CPs, the predicted fibronectin type III-like domain overlapped with the β sheet and loop dominated R domains that structurally resembled the R domains of human cathepsin B and cruzain (**Figure 3A**) - where heparin has been implicated to bind ^[57] [41]^. The negatively charged heparin occupied neutral-positive electrostatic patches in most of the different *NA* CPs (**Figure 3C**) at about the same location as the low-energy heparin-docked region of human cathepsin B ^[57]^. Similar results were obtained in this study despite adopting a different methodology of *sequence* pattern-based prediction and *structural* residue-profile derivation, to specify the region for non-covalently docking heparin, as compared to the Costa *et al* study, where they relied entirely on protein patches having positive electrostatic potential. Heparin bound to the *NA*-CPs, away from the active site, making the K and R contacts, as made by human cathepsin B’s K157 and R234 (**Figure 1**).

Heparin-like glycosaminoglycan present in animal plasma membrane and ECM, form components of the host-pathogen interface ^[45]^. The GAGs serve as recognition factors for molecular interactions, controlling activities like cell adhesion and parasitic infection ^[41] [45] [62]^. These negatively charged molecules are covalently linked to syndecans and glypicans proteoglycans of host cells, and help to recognize and bind parasitic cysteine proteases ^[45]^ such as cruzain ^[41] [62]^. The simultaneous binding of the proteases with their protein substrate and the latter’s covalently linked GAG results in the formation of ternary complexes ^[41] [62]^, which facilitates the enzymatic action of the proteases.

Heparin/heparin-sulfate binding to human cathepsin B protects the enzyme from alkaline pH induced loss of catalytic activity ^[57] [63]^. Inactivation of cathepsin B under alkaline conditions occurs due to disruption of electrostatic interactions like the thiolate-imidazolium ion pair of the active site, and breakage of crucial salt bridge interactions. Heparin binding prevents the loss of such interactions and helps stabilize the protease’s structure at alkaline pH by maintaining the helical content of the active enzyme ^[63]^. This allosteric mechanism mediated by heparin’s binding away from the active site has been corroborated by computational studies ^[57]^. *NA*-CPs being cathepsin B-like are likely to be affected by heparin binding in a similar way. The molecule docked on to the *NA*-CPs, at a region similar to what has been reported by Costa *et al* in their cathepsin B–heparin dockings (**Figure 1**). This is suggestive of possible allosteric control that could be exercised by heparin over the enzymatic action of the *NA*-CPs to make them functional even at alkaline pH, which the parasite encounters in blood during its migratory route to the host gut. Such controls by heparin could be crucial for the survival of hookworm, which encounters different pH conditions within the host, and whose cysteine proteases have varying degrees of overall surface electrostatics (**Figure 3C**).

### 3.5 Scope for heparin-based drug design

Heparin-reversal agents could counter the suggested blood-thinning activity by indigenous worm heparin. Anti-heparin agents had been previously designed to prevent excessive bleeding caused by clinically used heparin. ^[64]^. The molecular content of worm heparin, if and when characterized, could enable the design of appropriate positively charged anti-heparin molecules. Deglycosylation of the *NA* CPs that are covalently bound to GAGs, aided by enzymes found at the human small intestine ^[65]^ (where adult hookworms attach), could release free heparin at the extracellular matrix. Reversing the *worm-heparin-induced* (amongst other factors) anticoagulation can possibly prevent overt gastrointestinal bleeding caused by hookworm ^[66]^, and help thwart the parasite’s survival (**Figure 4**). *NA*-CPs’ suggestive role in hemoglobin degradation and anticoagulation by HMWK-cleavage, assisted by covalently bound heparin on host proteins, underscores the importance of abolishing non-covalent GAG interaction with the *NA*-CPs, for controlling the proteases’ pathogenic activities (**Figure 4**). Soluble heparin/heparin sulfate usage as competitive inhibitors have the chances of causing adverse physiological effects due to their anticoagulant/immunogenic properties ^[45]^. However, synthetic GAG mimetics with limited biological activity (minimum active structure) or polysaccharides ^[67] [68]^ targeted at GAG binding site (**Table 2**), could be used for inhibiting hookworm infection, in conjunction with active-site inhibitors.

**Figure 4.**
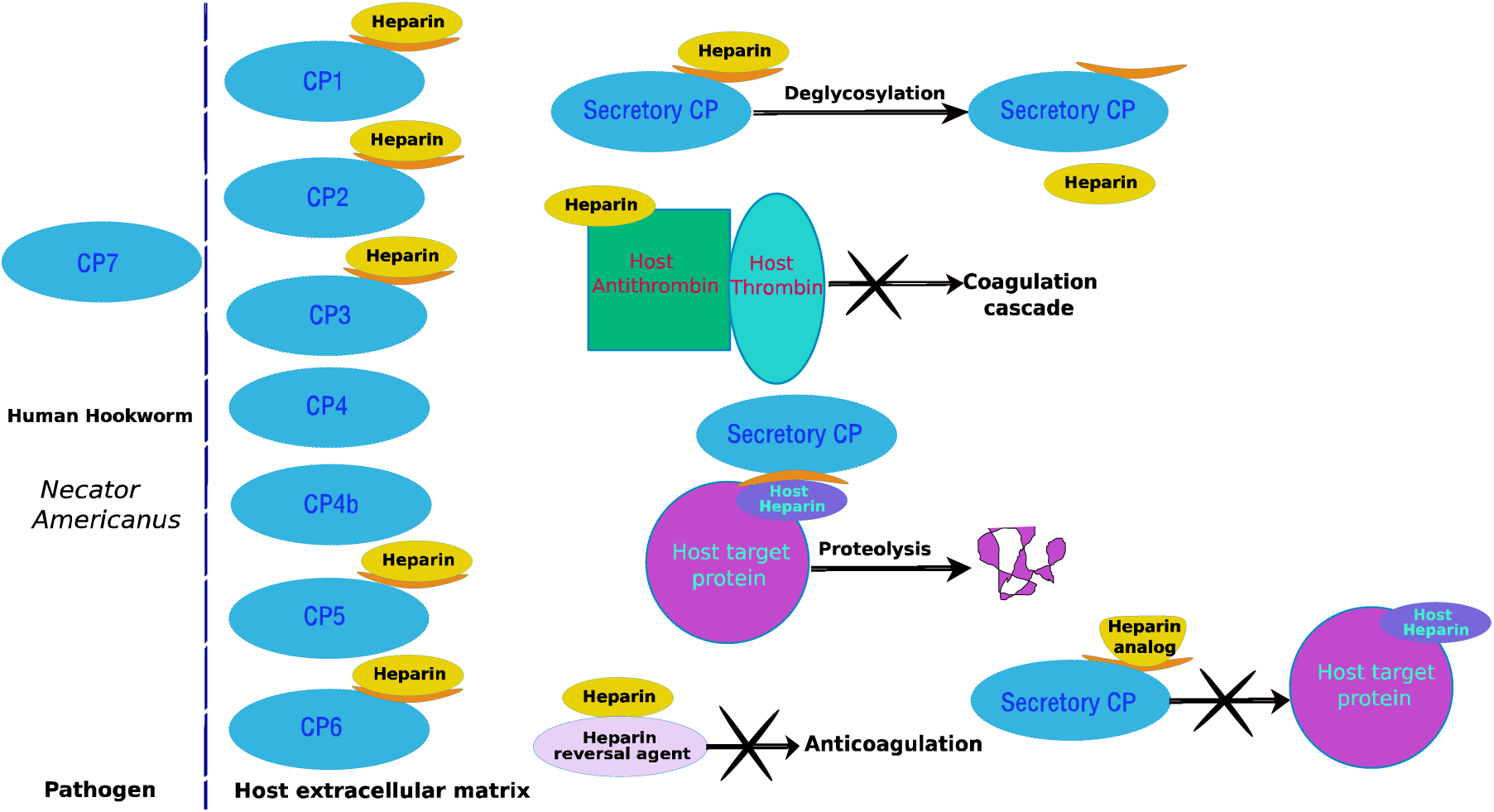
Schematic diagram summarizing the suggested *heparin-based* molecular interactions at the host-pathogen interface. The secretory CPs are on the extracellular matrix side, with their heparin-binding site shown in orange.

## 4 Conclusions

This study was initiated to explore the role of *NA* CPs in human hookworm pathogenesis, as the proteases mostly remain uncharacterized. Bioinformatics based analyses of the *NA*-CPs is indicative of the presence of CP1, CP2, CP3, CP4, CP4b, CP5 and CP6 in the ES content of the hookworm for host-pathogen interactions. The secretory proteases could be transporting worm heparin-like molecules to the extracellular matrix of the host for impairing coagulation, thereby suggesting the use of designed anti-heparin therapeutic molecules. The hemoglobinase motif derived here is harbored by CP2 and CP3, which suggest hemoglobinase activity in these two proteins, of which only CP3 has been confirmed to degrade globin ^[18]^. Possible HMWK – cleaving activity by *NA* CP1 and CP6 hint towards these proteases’ role in evading the host hemostatic system for preventing blood clots, in order to facilitate migration. The docking studies are speculative of *NA* CPs’ heparin-assisted HMWK-cleaving activity. Such assistance by heparin, already verified for the proteases from other parasites viz: *T.Cruzi* ^[40] [41]^ and *S.mansoni* ^[52]^ that navigate blood capillaries points to similar survival strategy adoption by larvae-stage *NA*, which traverses blood capillaries. This study is an attempt to provide molecular level information based on computational predictions, decoding previously unreported functions of the individual *NA* cysteine proteases and the role that heparin could be playing. The extracellular localizations of CP1-CP6 and the delineated pathogenic roles of CP1, CP2, CP3 and CP6, render these CPs to be multi-targeted by designed heparin analog, which could help inhibit hookworm infection, alongside active-site inhibitors.

## Acknowledgement

Self-funding by the first author is acknowledged. The authors thank useful discussions with Prof. Conor Caffrey, UCSD. The master’s dissertation work on hookworm by Kevin Weidmer, from which only the CP sequence IDs were noted is also acknowledged.

## References

[1] P.J. Hotez, S. Brooker, J.M Bethony, M.E. Bottazzi, A. Loukas, S.H Xiao, N Engl J Med. 2004, 351:799–807.

[2] P.J. Hotez, J.M. Bethony, D.J. Diemert, M. Pearson, A. Loukas, Nat. Rev. Microbiol. 2010, 8:814–826

[3] A. Loukas, S. Gaze, J.P. Mulvenna, R.B. Gasser, P.J. Brindley, D.L. Doolan, J.M. Bethony, M.K. Jones, G.N. Gobert, P. Driguez, D.P. McManus, and P.J Hotez, OMICS 2011 15(9): 567–577

[4] D.J. Diemert, J.M. Bethony, P.J Hotez, Clin Infect Dis. 2008, 46: 282–288.

[5] M.S. Pearson, L. Tribolet, C. Cantacessi, M.V. Periago, M.A. Valerio, A.R. Jariwala, P.J. Hotez, D. Diemert, A. Loukas, J. Bethony, J allergy clin immunol. 2012, vol 130, No.1

[6] N.J. Kassebaum, R. Jasrasaria, M. Naghavi, S.K. Wulf, N. Johns, R. Lozano, M. Regan, D. Weatherall, D.P. Chou, T.P. Eisele, S.R Flaxman, R.L Pullan, S.J. Brooke, C.J. Murray, Blood. 2014. 123:615–624.

[7] H. Sakti, C. Nokes, W.S. Hertanto, S. Hendratno, A. Hall, D.A. Bundy, Satoto, Trop Med Int Health. 1999, 4(5): 322–34.

[8] P. Hotez, Ann NY Acad Sci. 2008, 1136: 38–44

[9] J.C. Vetter, M.E. van der Linden Z Parasitenkd. 1977, 53:255–62.

[10] J.C. Vetter, M.E. van der Linden Z Parasitenkd. 1977, 53:263–66.

[11] S. Brooker, J. Bethony, P.J. Hotez, Adv Parasitol. 2004, 58: 197–288.

[12] A. Brown, J.M. Burleigh, E.E. Billett, D.I. Pritchard, Parasitology. 1995, 110: 555–563.

[13] R. Bungiro, M. Cappello, Curr Infect Dis Rep 2011 13:210–7.

[14] N. Ranjit, M.K. Jones, D.J. Stenzel, R.B. Gasser, A. Loukas, Int J for Parasitol 2006 36: 701–710.

[15] D.P. Jasmer, J. Roth, P.J. Myler. Mol. Biochem. Parasitol. 2001, 116:159–169.

[16] M. Sajid, J.H. McKerrow, Mol. Biochem Parasitol. 2002, 120: 1–21

[17] V. Turk, V. Stoka, O. Vasiljeva, M. Renko, T. Sun, B. Turk, D. Turk, Biochimica et Biophysica Acta. 2012, 68–88.

[18] N. Ranjit, B. Zhan, B. Hamilton, D. Stenzel, J. Lowther, M. Pearson, J. Gorman, P. Hotez, A. Loukas, J Infect Dis. 2009, 199:904–12.

[19] P. Stanssens, P.W. Bergumt, Y. Gansemanst, L. Jesperst, Y. Larochet, S. Huangt, S. Makit, J. Messenst, M. Lauwereyst, M. Cappello, P.J. Hotez, I. Lasterst, G.P. Vlasuk, Proc. Natl. Acad. Sci. USA. 1996, Vol. 93: 2149–2154.

[20] L.M. Harrison, A. Nerlinger, R.D. Bungiro, J.L. Cordova, P. Kuzmic, M. Cappello, J Biol Chem. 2002 Vol. 277(8), Issue of February 22: 6223–6229.

[21] B.A. Furmidge, L.A. Horn, D.I. Pritchard, Parasitol. 1995, 112: 81–87.

[22] G.N. Cox, D. Pratt, R. Hageman, R.J. Goisvenue, Mol Biochem Parasitol. 1990, 41: 25–34.

[23] N. Ranjit, B. Zhan, D.J. Stenzel, J. Mulvenna, R. Fujiwara, P.J. Hotez, A. Loukas, Mol Bioc/hem Parasitol 2008. 160: 90–99.

[24] *The UniProt Consortium*. 2015. Vol. 43, Database issue.

[25] R.A. Abagyan, M.M. Totrov, D.A. Kuznetsov, J. Comp. Chem. 1994, 15, 488–506.

[26] G. Blobel, PNAS. 1980, Mar; 77(3): 1496–1500.

[27] I. Jonassen, J.F. Collins, D. Higgins, Prot Sci. 1995, 4(8): 1587–1595.

[28] C.J.A. Sigrist, L. Cerutti, N. Hulo, A. Gattiker, L. Falquet, M. Pagni, A. Bairoch, P. Bucher, Brief Bioinform. 2002, 3:265–274.

[29] T.N. Petersen, S. Brunak, G.V. Heijne, H. Nielsen, Nat Methods. 2011, 8:785–786.

[30] O. Emanuelsson, S. Brunak, G.V. Heijne, H. Nielsen, Nat Protocols. 2007, 2: 953–971.

[31] H. Bannai, Y. Tamada, O. Maruyama, K. Nakai, S. Miyano, Bioinform. 2002,18(2): 298–305.

[32] A. Krogh, B. Larsson, G.V. Heijne, E.L. Sonnhammer, J Mol Biol. 2001, Jan 19; 305(3): 567–80.

[33] B.R. King, C. Guda, Gen Biol. 2007, 8(5): R68

[34] A. Pierleoni, P.L. Martelli, P. Fariselli, R. Casadio, Bioinform. 2006 22 (14): e408–e416.

[35] M. Bodén, J. Hawkins, 2005. Bioinform. 21(10): 2279–2286.

[36] C.S. Yu, Y.C. Chen, C.H. Lu, J.K. Hwang, Prot: Str Funct Bioinform. 2006. 64:643–651.

[37] K. Hiller, A. Grote, M. Scheer, R. Münch, D. Jahn, Nucl Acids Res. 2004.32: W375–W379.

[38] D.A. Benson, M. Cavanaugh, K. Clark, I. Karsch-Mizrachi, D.J. Lipman, J. Ostell, E.W. Sayers, Nucl Acids Res. 2013, Jan; 41 (Database issue): D36–42.

[39] N.M.T. Barros, I.L. Tersariol, M.L. Oliva, M.S. Araujo, C.A. Sampaio, L. Juliano, G. Motta, Biol. Chem. 2004. 385: 551–555

[40] E. Del Nery, M.A. Juliano, A.P. Lima, J. Scharfstein, L. Juliano, J Biol Chem 1997 Vol. 272(41): 25713–25718.

[41] A.P.C.A. Lima, P.C. Almeida, I.L.S. Tersariol, V. Schmitz, A.H. Schmaier, L. Juliano, I.Y. Hirata, W. Muller-Esterl, J.R. Chagas, J. Scharfstein, JBC. 2002, Vol. 277, No. 8, Issue of Feb 22: 5875–5881.

[42] M. Maurer, M. Bader, M. Bas, F. Bossi, M. Cicardi, M. Cugno, P. Howarth, A. Kaplan, G. Kojda, F. Leeb-Lundberg, J. Lötvall, M. Magerl,. Allergy. 2011, 66: 1397–1406.

[43] H.M. Berman, J. Westbrook, Z. Feng, G. Gilliland, T.N. Bhat, H. Weissig, I.N. Shindyalov, P.E. Bourne, Nucl Acids Res, 2000 28: 235–242.

[44] S. Gillmor, C.S. Craik, R.J. Fletterick, Prot Sci. 1997, 6:1603–1611.

[45] A.H. Bartlett, P.W. Park, Exprt Rev Mol Med. 2010, 1–25.

[46] C. Bru, E. Courcelle, S. Carrère, Y. Beausse, S. Dalmar, D. Kahn, Nucl Acids Res. 2005, 33: D212–D215.

[47] C. Illy, O. Quraishi, J. Wang, E. Purisima, T. Vernet, J.S. Mort, J. Biol. Chem. 1997, 272: 1197–1202.

[48] R.A. Laskowski, M.W. MacArthur, D.S. Moss, J. Appl. Cryst, 1993, 26: 283–291.

[49] R. Pankov, K.M. Yamada. J. Cell Sci. 2002 115: 3861–3863.

[50] S. Baig, R.T. Damian, D. S. Peterson, Exptl Parasitol. 2002, 101: 83–89.

[51] M.M. Mebius, P.J.J. van Genderen, R.T. Urbanus, A.G.M. Tielens, de P.G. Groot, PLoS Pathog. 2013, 9(12): e1003781.

[52] W.S. Carvalho, C.T. Lopes, L. Juliano, P.M. Coelho, J.R. Cunha-Melo, W.T. Beraldo, J.L. Pesquero, Parasitol. 1998, 117: 311–319.

[53] A. Da’dara, P.J. Skelly, Blood Rev. 2011, 25: 175–179.

[54] L. Herszenyi, M. Plebani, P. Carraro, M. De Paoli, R. Cardin, F. Di Mario, S. Kusstatscher, R. Naccarato, F. Farinati, Am j. gastroenterology. 1997, 92: 843–847

[55] F. Fenaille, M.L. Mignon, C. Groseil, C. Ramon, S. Riandé, L. Siret, N. Bihoreau, Glycobiol. 2007, Vol. 17 (9): 932–944.

[56] S. Takahashi, M. Leiss, M. Moser, M. Ohashi, T. Kitao, D. Heckmann, A. Pfeifer, H. Kessler, J. Takagi, H.P. Erickson, R. Fässler, J Cell Biol, 2007. Vol 178(1): 167–178.

[57] M.G.S Costa, P.R. Batista, C.S. Shida, C.H. Robert, P.M. Bisch, P.G. Pascutti, BMC Genomics. 2010. 11(Suppl 5): S5.

[58] M. Forster, B. Mulloy, Biochem Soc Trans. 2006, 34:431–434.

[59] A.E.I. Proudfoot, S. Fritchley, F. Borlat, J.P. Shaw, F. Vilbois, C. Zwahlen, A. Trkola, D. Marchant, P.R. Clapham, T.N.C. Wells, J Biol Chem. 2001, 276: 10620–10626.

[60] D.M. Mann, E. Romm, M. Migliorini, J Biol Chem. 1994, 269:23661–23667.

[61] J.R. Fromm, R.E. Hileman, E.E.O. Caldwell, J.M. Weiler, R.J. Linhardt, Arch Biochem Biophys. 1997, 343:92–100.

[62] W.A.S. Judice, M.A. Manfredi, G.P. Souza, T.M. Sansevero, P.C. Almeida, C.S. Shida, T.F. Gesteira, L. Juliano, G.D. Westrop, S.J. Sanderson, G.H. Coombs, I.L.S. Tersariol, PLoS ONE. 2013, 8(11): e80153.

[63] P.C. Almeida, I.L. Nantes, J.R. Chagas, C.C.A. Rizzi, A. Faljoni-Alario, E. Carmona, L. Juliano, H.B. Nader, I.L.S. Tersariol, J.Biol.Chem 2001. Vol. 276, No. 2, Issue of Jan 12: 944–951.

[64] R. A. Shenoi, M.T. Kalathottukaren, R. J. Travers, B. F. L. Lai, A. L. Creagh, D. Lange, K. Yu, M. Weinhart, B. H. Chew, C. Du, D. E. Brooks, C.J. Carter, J. H. Morrissey, C. A. Haynes and J. N. Kizhakkedathu, Sci Trans Med. 2014. Vol. 6, Issue 260, pp. 260ra150.

[65] A.J. Daya, M. S. DuPonta, S. Ridleyb, M. Rhodesc, M.J.C. Rhodesa, M.R.A. Morgana, G. Williamson, FEBS Letters. 1998. 436: 71–75

[66] J.M. Chen, X. M. Zhang, L. J. Wang, Y. Chen, Q. Du, J.T. Cai, Asian Pacific J. Trop Med. 2012. 331–332.

[67] W.F. Vann, M.A. Schmidt, B. Jann, K. Jann, Eur. J. Biochem. 1981, 116: 359–364.

[68] R. Copeland, A. Balasubramaniam, V. Tiwari, F. Zhang, A. Bridges, R.J. Linhardt, D. Shukla, J. Liu, Biochem. 2008 47:5774–5783.

