## supplementary material for "Cysteine proteases of human hookworm *Necator Americanus as virulence* factors and implications for drug design with anti-heparin and heparin analogs: A bioinformatics study"

| Protease | Residue: atom | Residue: atom | Distance in Armstrong |
| --- | --- | --- | --- |
| Cruzain | Ala137: CA | Glu207: CA | 5.47 |
| Human Cathepsin B | Ala172: CA | Glu244: CA | 5.58 |
| CP1 | Ala175: CA | Asp246: CA | 7.75 |
| **CP6** | **Ala174: CA** | **Glu245: CA** | **7.79** |

**Supplementary Table 1.** Distance in angstrom between the C-alpha atoms of the critical Asp/Glu and the conserved Ala of the substrate cleft, measured from the crystal structures/homology models of the HMWK-cleaving enzymes. The residue numberings pertain to the mature proteases.

| *NA* Cysteine Protease | Sequence identity percent | Sequence similarity percent |
| --- | --- | --- |
| CP5 | 100 | 100 |
| CP1 | 52.7 | 75.3 |
| CP2 | 55.26 | 78.64 |
| CP3 | 53.16 | 75.23 |
| CP4 | 50.64 | 65.11 |
| CP4b | 50.64 | 65.11 |
| CP6 | 54.05 | 74.97 |
| CP7 | 40.74 | 55.15 |

**Supplementary Table 2.** Pairwise sequence identity and similarity of the *NA* CPs with respect to CP5, within the fibronectin domain alignment region.
